## Supplementary Information for "Higher social tolerance is associated with more complex facial behavior in macaques"

Alan V. Rincon

Jérôme Micheletta

**This PDF file includes:**

Supplementary text

Tables S1 to S2

Figure S1

### Study subjects and data collection

Behavioral data and video recordings were collected on one adult male and 31 adult female rhesus macaques (*M. mulatta*), on 18 adult male and 28 adult female Barbary macaques (*M. sylvanus*), and 17 adult male and 21 adult female crested macaques (*M. nigra*). Admittedly, a more balanced sample size per sex would have been preferable for rhesus macaques. Nevertheless, male and female macaques must (and do) interact and communicate with each other regularly. Therefore, we have no *a priori* reason to expect an overall difference in the diversity and complexity of facial behavior between the sexes. The social complexity hypothesis makes predictions at the level of societies, and we feel like our sample size for rhesus macaques is large enough to representatively capture the complexity of their facial behavior.

Rhesus macaques belonged to one breeding group (Gruppe 1) at the German Primate Center, Germany. Monkeys were housed in naturalistic outdoor enclosure (approximately 290m$^2$ and 4-7m high) with free access to a heated indoor area (approximately 80m$^2$ and 5-7m high), which were enriched with ropes, logs, swings, and a small pond. Monkeys were fed daily a variety of fruits and vegetables, nuts, seeds, cereals, commercial monkey pellets, and had *ad libitum* access to water. All observations, including the recording of videos, were conducted outside of the enclosures. Data collection on the rhesus macaques took place between June and October 2021. Barbary macaques belonged to one group (German Group) out of two groups living at Trentham Monkey Forest, United Kingdom. Monkeys were able to freely move within a 24-hectare open enclosure of forest and grassy areas. Monkeys were fed daily a variety of fruits, vegetables, seeds, and monkey chow, and had *ad libitum* access to water. Data collection on the Barbary macaques took place between August and November 2019. Crested macaques belonged to two wild groups (R2A and PB1B) living in Tangkoko-Batuangus Nature reserve, North Sulawesi, Indonesia, and observed within the Macaca Nigra Project (http://www.macaca-nigra.org). Monkeys were not provisioned by humans and fed on natural foods and were habituated to the presence of human observers. Data collection on the crested macaques took place between December 2018 and April 2019.

### Random forest classifier predictions

**Table S1:** Confusion matrices for random forest classifier predictions of social context from Action Unit combinations.

| | Truth | | |
|---|---|---|---|
| Prediction | affiliative | aggressive | submissive |
| rhesus | | | |
| affiliative | 636 | 19 | 9 |
| aggressive | 81 | 1,205 | 17 |
| submissive | 2 | 6 | 731 |
| Barbary | | | |
| affiliative | 2,573 | 24 | 442 |
| aggressive | 200 | 1,219 | 165 |
| submissive | 166 | 34 | 528 |
| crested | | | |
| affiliative | 1,134 | 90 | 43 |
| aggressive | 16 | 86 | 11 |
| submissive | 3 | 1 | 7 |

### Action Unit list

**Table S2:** Action Units (AU) and Descriptors (AD) observed and coded in the study.

| Action Unit/Descriptor | Description |
|---|---|
| AU1+2 | Brow raiser |
| AU41 | Glabella (brow) lowerer |
| AU5 | Upper lid raiser |
| AU6 | Cheek raiser |
| AU8 | Lips toward each other |
| AU9 | Nose wrinkler |
| AU10 | Upper lip raiser |
| AU12 | Lip corner puller |
| AU16 | Lower lip depressor |
| AU17 | Chin raiser |
| AU18* | Lip pucker |
| AU25 | Lips parted |
| AU26 | Jaw drop |
| AU27 | Jaw stretch |
| AU43 | Eyelid droop |
| EAU1 | Ears forward |
| EAU2 | Ears elevator |
| EAU3 | Ears flattener |
| AD19 | Tongue show |
| AD29 | Jaw thrust |
| AD59 | Head toss |
| AD101 | Scalp retraction |
| AD181† | Lipsmack |
| AD182† | Teeth chatter |
| AD184† | Jaw wobble |
| AD185‡ | Jaw oscillation |

*AU18i and AU18ii were combined because it was difficult to reliably distinguish between them when coding videos. †Excluded in favor of AD185, which allows for a more detailed description of facial behavior when combined with other AUs. ‡New AD not previously described in MaqFACS. Denotes stereotyped up-and-down movement of the jaw.

### Specificity bias correction

Facial Action Coding System (FACS) data were simulated for three contexts (A, B, C) and 10 elements (1 to 10, representing Action Units). Specificity was calculated when all contexts had an equal number of observations (denoting the true specificity) and on a subset of the data where the number of observations between the three contexts was imbalanced at a ratio of 10:5:1. Specificity values were skewed higher in the context with most observations (A) and skewed lower in context with fewest observations (C). Upsampling the minority contexts, such that all contexts had the same number of observations, substantially minimized the error bias in specificity values (Figure S1). The R script for the simulation can be found at https://github.com/avrincon/macaque-facial-complexity.

**Figure S1:** Calculating context specificity on an imbalanced dataset. Specificity was calculated on a simulated dataset with an imbalanced number of observations per context. The calculated specificity values deviated from the true specificity such that they were higher in the context with most observations and lower in the context with fewest observations (green dots). Randomly upsampling observations from the minority contexts (B and C) such that they have the same number of observations as the majority context (A) prior to calculating specificity minimized the bias in the calculated specificity values (purple dots)

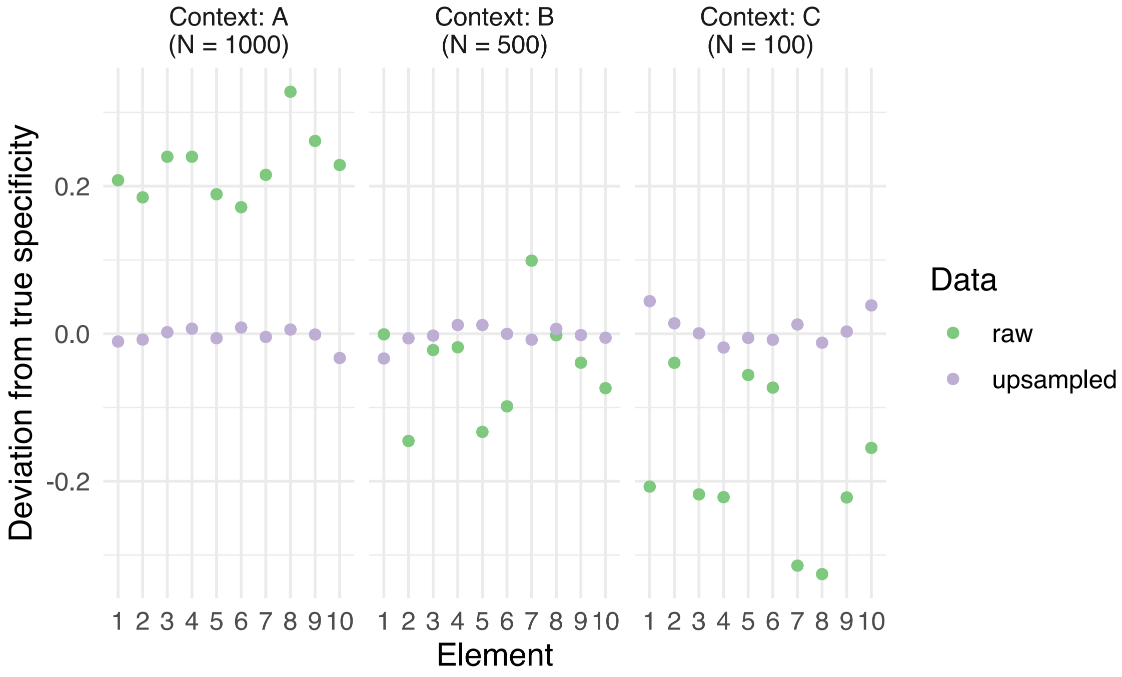
